# Brahmi Ghrita Exerts Nephroprotective Effects by Restoring Cytoskeletal Integrity and Ion Transport in *Drosophila* Model of Polycystic Kidney Disease

**DOI:** 10.64898/2026.02.13.705756

**Authors:** Saurabh Chand Sagar, Madhu G Tapadia

## Abstract

**Background:** Polycystic kidney disease (PKD) is a genetic disorder characterized by progressive cyst formation, epithelial disorganization, and impaired fluid transport, ultimately leading to renal failure. Disruption of cytoskeletal dynamics and epithelial polarity is central to PKD pathogenesis. Malpighian tubules (MTs) of *Drosophila melanogaster* serve as a conserved renal analog, and caspase-3/Drice–deficient flies exhibit a robust PKD-like tubular phenotype, providing a powerful in vivo model to investigate therapeutic interventions.

**Purpose:** This study evaluates the therapeutic potential of the Brahmi Ghrita (BG) in ameliorating PKD-like defects in *Drosophila* Drice mutants and elucidates the underlying cellular and molecular mechanisms.

**Methods:** Drice mutant flies were reared with dietary BG supplementation, and developmental viability (pupation and eclosion) was assessed. Tubule morphology was analyzed by measuring cyst formation and tubule dimensions. Stellate cell (SC) number, shape, and nuclear size were quantified. Cytoskeletal organization and epithelial polarity were examined using F-actin and polarity markers. Molecular analyses included assessment of Rho1 signaling, Gelsolin, and Rho kinase (Rok) localization. Tubule physiology was evaluated by uric acid crystal deposition and Na□/K□-ATPase expression.

**Results:** BG supplementation significantly improved pupation and eclosion rates in Drice mutants and markedly reduced cystic dilation by restoring tubule width without altering developmental length. BG selectively increased stellate cell number and normalized aberrant morphology, while principal cell number remained unchanged. Cytoskeletal disorganization and polarity defects were rescued, accompanied by normalization of elevated Rho1 levels and restoration of the actin-severing protein Gelsolin. BG enhanced Na□/K□-ATPase expression and reduced uric acid accumulation, consistent with improved epithelial transport function. Additionally, BG promoted nuclear enrichment of Rok, indicating altered Rho-associated signaling dynamics.

**Conclusion:** Brahmi Ghrita confers nephroprotective effects in a genetic PKD model by coordinately restoring cytoskeletal integrity, epithelial polarity, and ion transport machinery. Rather than broadly suppressing Rho signaling, BG appears to rebalance the Rho1–Gelsolin axis and re-establish transport competency, culminating in structural and functional rescue. These findings provide mechanistic evidence supporting BG as a multi-target modulator of epithelial homeostasis in PKD-relevant contexts.

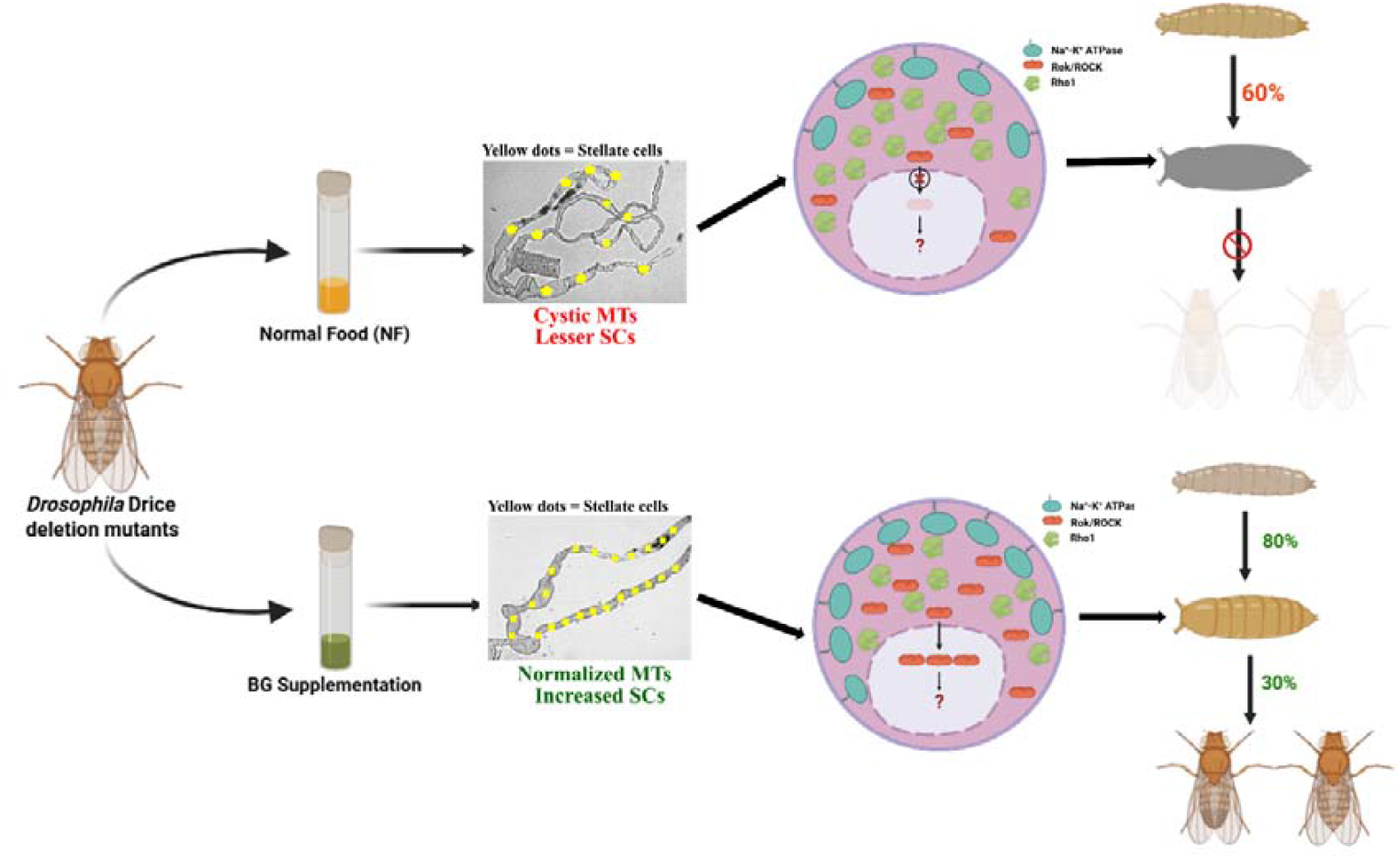

## 1. Introduction

Polycystic kidney disease (PKD) is characterized by the formation of multiple fluid-filled cysts within the renal system [1]. The cysts typically emerges as a dilation of renal epithelial tubules, in the ureteric region of the kidney, which progressively expands to involve the entire organ [2] [3]. The distal segments of the nephrons and collecting ducts are often the most severely affected. As the disease advances, cystic lesions can also develop in other epithelial organs, such as the liver and pancreas, ultimately leading to end-stage renal disease [2] [3]. The hallmark features of PKD include abnormal proliferation of renal epithelial cells, formation of multiple fluid-filled cysts, disruption of tissue morphology, extracellular matrix organization, and cellular differentiation [4]. Malpighian tubules (MTs) in *Drosophila* are analogous to the human kidneys, regulating excretion, maintaining osmoregulation and exhibiting immune response [5, 6]. MTs in *Drosophila* are long thin paired structure joined to the junction of midgut and hindgut via short ureter region. One pair project anteriorly toward the midgut and is referred to as the anterior tubules (ATs), whereas the other pair extends posteriorly toward the hindgut and is termed the posterior tubules (PTs) (Wessing and Eichelberg, 1978). MTs of *Drosophila melanogaster* comprise two major cell types, namely the principal cells (PCs) and stellate cells (SCs) [7]. PCs are the predominant cell type, ectodermal in origin, whereas SCs are lesser in number as well as size, mesodermal in origin, and interspersed among PCs along the length of the tubule [8]. MTs in *Drosophila* does not undergo histolysis/cell death despite showing the caspase activity in them [9, 10]. MTs in Caspase-3/Drice-deficient *Drosophila* exhibit polycystic phenotype that closely mirrors PKD-associated defects in human kidneys [10] [11]. Drice deficiency leads to morphological abnormalities linked to altered Rho1 GTPase signaling, as well as physiological impairments that compromise tubule function [10, 11]. The striking resemblance between these morphological and physiological defects in Drice-deficient MTs and those characteristic of PKD in vertebrates supports *Drosophila* as an excellent model to investigate the molecular mechanisms underlying PKD [10].

Ayurveda, the traditional system of medicine, originated during the Vedic period; is native to the Indian subcontinent [12]. Ayurvedic formulations are categorized into eight major branches, among which Rasayana Tantra focuses on rejuvenation, longevity, and the enhancement of intellect and physical strength. BG is a well-known Ayurvedic Rasayana belonging to the Medhya Rasayana group of Ayurveda. BG is primarily recognized as a brain tonic with multiple therapeutic benefits, including improvement of learning and memory [13], reduction of amnesia [14], and anticonvulsant activity [15]. Recent studies have demonstrated its protective role against morphine-induced nephrotoxicity in rats [16]. Considering the broad therapeutic potential of BG, particularly its recently reported nephroprotective properties, we aim to investigate its effects on cellular architecture and excretory function in Caspase 3 deletion mutation in *Drosophila viz*., *Drice*^*Δ2C8*^*/Drice*^*Δ2C8*^.

In this study, we explored the therapeutic aspects of BG. BG supplementation markedly enhanced the survival of *Drice* mutant progeny, enabling a higher proportion of flies to successfully eclose into adults. Notably, BG feeding significantly reduced cyst formation and mitigated the excessive dilation of MTs observed in the mutant background, restoring their morphology closer to that of the wild type. At the cellular level, BG treatment alleviated nuclear enlargement and increased the number of SCs in the MTs of *Drice* mutants, indicating a substantial rescue of tubular secretion. Further mechanistic analyses revealed that BG feeding restored actin organization and cell polarity in the Drice mutant MTs, resembling wild-type characteristics. Elevated Rho1 expression in *Drice* mutants was significantly reduced following BG administration. BG treatment also, decreased uric acid crystal accumulation and improves fluid secretion rates in the mutant flies, indicating a concomitant recovery of physiological function. Collectively, our results highlight the nephroprotective potential of BG and demonstrate its ability to partially rescue the PKD-like phenotype in *Drosophila*.

## 2. Results

### 2.1: Brahmi Ghrita reduces larval & pupal mortality and improves eclosion rate in Drice mutants

BG supplementation has been reported to exert a nephroprotective effect against morphine-induced toxicity in mice [16], we aimed to investigate the effect of BG supplementation in *Drosophila* Drice homozygous mutants. In order to determine the appropriate dose concentration for the drug administration, we fed wild type *Drosophila* on the BG supplementation (BG 0.1%, BG 0.25%, BG 0.50% and BG 1.0% supplementation), to determine the LC50 dose and its effect on the fecundity. BG supplementation altered the median life span of 38 days for flies fed on normal food (NF), to the 36 days, 35 days, 30 days and 26 days for the BG 0.10%, BG 0.25%, BG 0.50% and BG 1.0% supplementation respectively (Supplementary Table S1). Kaplan–Meier survival analysis revealed a non-significant decrease in the life span upon BG 0.10% and BG 0.25% supplementation concentration, these two dose along with one higher dose BG 0.50% were selected for further studies (Figure 1,A). Next, we examined the fecundity pattern upon BG 0.10%, BG 0.25% and BG 0.50% supplementation. BG 0.10% and BG 0.25% supplementation do not show significant change in the fecundity pattern; however, BG 0.50% supplementation reduced the fecundity significantly (Figure 1, B). Based upon these observation BG 0.25% was selected as optimum dose for drug administration with BG 0.50% selected as one higher concentration. To evaluate the efficacy of BG feeding on Drice mutants, BG 0.25% was selected as the optimum doses, while BG 0.50% and BG 1.0% were included as two higher concentrations for the study.

**Figure 1:**
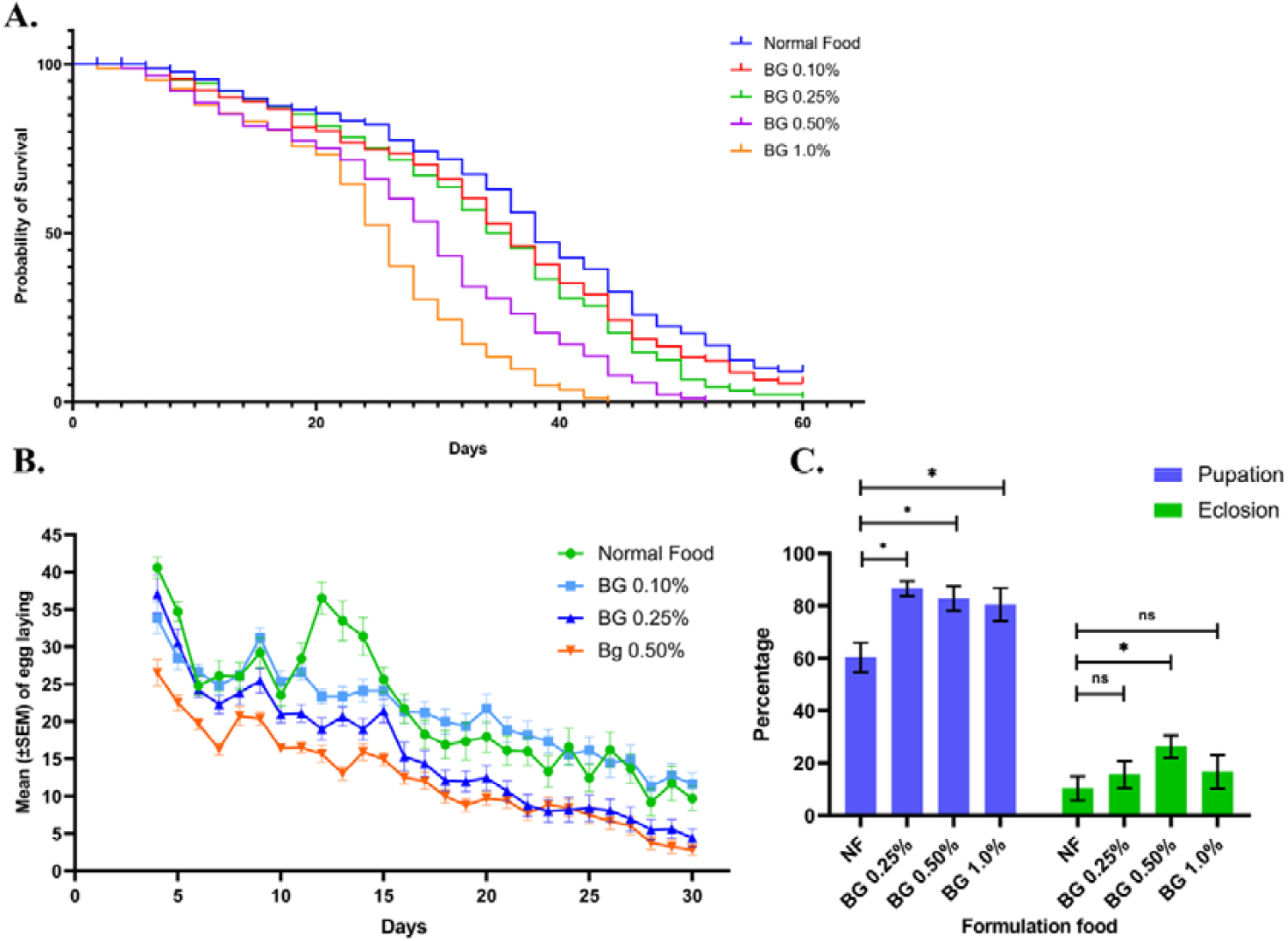
Effect of BG supplementation on fly survival. **(A) Longevity in wild-type Drosophila.** BG supplementation at 0.10% and 0.25% did not adversely affect normal lifespan, whereas, BG 0.50% and BG 1.0% supplementation reduces the median life span of the flies. The median lifespan of flies fed on NF was 38 days, which changed to 36, 35, 30, and 26 days upon supplementation with BG 0.10%, 0.25%, 0.50%, and 1.0%, respectively (n = 500). Statistical significance was determined using survival analysis followed by log-rank test. **(B) Fecundity in wild-type Drosophila**. BG 0.10% supplementation showed no effect on fecundity (p = 0.9923), while BG 0.25% showed a marginal reduction (p = 0.482), and BG 0.50% significantly reduced fecundity (p = 0.002) (n = 20 pairs). Statistical significance was determined using Welch’s one-way ANOVA followed by Dunnett’s multiple comparison test. p < 0.05 was considered statistically significant (*p < 0.05). **(C) Survival in Drice mutant flies**. BG supplementation improved pupation at all tested concentrations—BG 0.25% (p = 0.0127), BG 0.50% (p = 0.0154), and BG 1.0% (p = 0.0358). For eclosion, BG 0.10% (p = 0.5249) and BG 1.0% (p = 0.4958) showed no significant effect, whereas BG 0.25% significantly enhanced eclosion (p = 0.0301) (n ≥ 300). Statistical significance was determined using Welch’s one-way ANOVA followed by Dunnett’s multiple comparison test. p < 0.05 was considered statistically significant (*p < 0.05).

Loss of Caspase-3 activity in *Drice* mutants leads to pronounced morphological impairments in the Malpighian tubules, which severely compromise organismal viability. Consequently, only 60–65% of larvae progress to the pupal stage, and approximately 10–12% of pupae successfully eclose as adults, with the emergent adults exhibiting markedly reduced survivability and dying shortly after eclosion. Upon feeding Drice mutants with different BG supplementation; pupation rate, which was 60.33 ± 5.6% in untreated mutants, increased to 86.67 ± 2.8%, 82.83 ± 4.7%, and 80.50 ± 6.3% for BG 0.25%, BG 0.50%, and BG 1.0%, respectively, indicating a significant improvement in pupation (Figure 1, C; Supplementary Table S2).

Similarly, the eclosion rate, which was 10.33 ± 4.6% in untreated Drice mutants, increased to 15.50 ± 5.4%, 26.33 ± 4.3%, and 16.67 ± 6.4% upon feeding BG 0.25%, BG 0.50%, and BG 1.0%, respectively, suggesting a notable enhancement in fly eclosion (Figure 1, C; Supplementary Table S2). The improvement was most pronounced at BG 0.50%, whereas BG 1.0% resulted in a marked reduction in eclosion, suggesting dose-dependent toxicity at higher concentrations. Therefore, BG 0.25% and BG 0.50% supplementations were selected for subsequent experiments on the Drice mutant flies.

### 2.2. BG supplementation restores tubule morphology in Drice mutants

Malpighian tubules (MTs) of Drice mutants displayed severely distorted morphology, characterized by multiple fluid-filled cysts, clustered cells and irregular twisting along their length, in contrast to the thin, elongated, and uniformly wide MTs of wild-type flies [10]. Upon BG supplementation, a marked improvement in tubule morphology was observed in Drice mutants reared on BG 0.25% (Figure 2, A-c) and BG 0.50% (Figure 2, A-d) supplementation when compared to those maintained on NF (Figure 2, A-b) (Refer to the Supplementary Figure S6 for tubule morphology at lower magnification). First, we checked the effect of BG on the length of the tubules as both, ATs and PTs of Drice mutants are significantly shorter than those of the wild type [10]. However, BG supplementation did not alter tubule length in either the ATs or PTs of Drice mutants (Figure 2, B; Supplementary Table 3).

**Figure 2:**
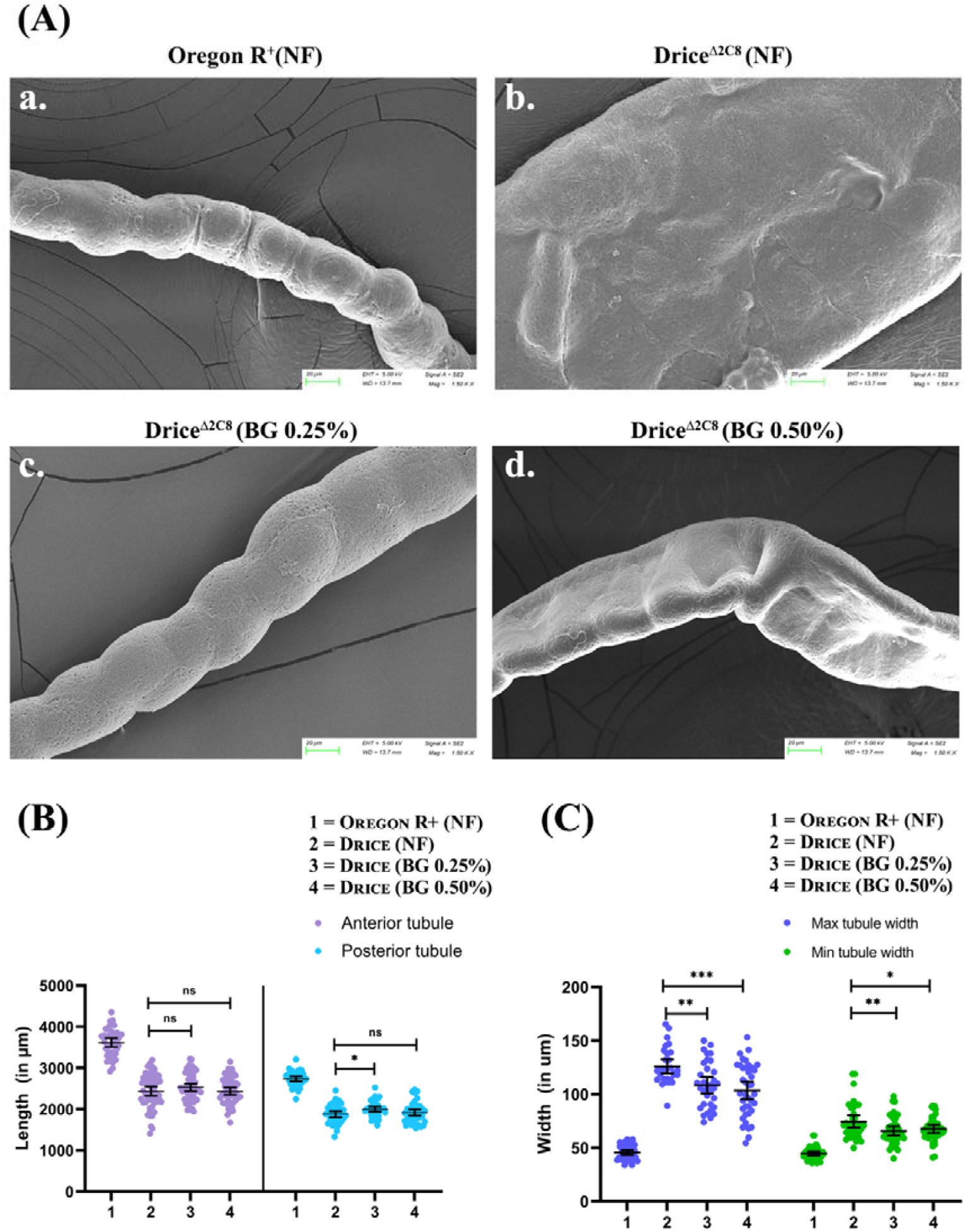
Restoration of tubule morphology in *Drice* mutants upon BG supplementation. **(A) Tubule morphology**. BG supplementation reduces tubule width significantly in BG 0.25% (c) and BG 0.50% (d) groups compared to *Drice* mutants on NF (b), while (a) shows wild-type morphology. Insets highlight selected regions. Scale bar = 100 µm. **(B) Tubule length**. ATs length remained unchanged in BG 0.25% (p = 0.4833) and BG 0.50% (p > 0.999) groups compared to *Drice* mutants fed on NF. However, a marginally significant increase was observed in BG 0.25% supplementation (p = 0.0422), while BG 0.50% showed no significant difference (p = 0.8627). Data are presented as Mean ± SEM (n ≥ 40). **(C) Reduction of cyst formation and tubule width in *Drice* mutants**. Tubule width in wild-type flies fed on NF (max = 46.67 ± 8.8 µm; min = 44.7 ± 6.25 µm) increased markedly in *Drice* mutants (max = 130.14 ± 22.3 µm; min = 74.26 ± 16.1 µm). BG supplementation significantly reduced tubule width in BG 0.25% (max = 108.91 ± 21.8 µm, p = 0.0020; min = 64.12 ± 11.2 µm, p = 0.0058) and BG 0.50% (max = 102.31 ± 23.7 µm, p = 0.0004; min = 65.98 ± 10.3 µm, p = 0.0145) groups, indicating morphological restoration in *Drice* mutants upon BG treatment (n ≥ 30 per group). Statistical significance was assessed using Welch’s one-way ANOVA followed by Dunnett’s multiple comparison test. p < 0.05 was considered statistically significant (*p < 0.05).

Given the cystic dilation observed in Drice mutants, we next examined the tubule width to assess the effect of BG feeding. To examine the tubule width, region with widest and narrowest tubule was measured. Both BG 0.25% and BG 0.50% supplementation significantly reduced tubule width compared to the flies fed on NF (Figure 2, C; Supplementary Table 4). Collectively, these results demonstrate that BG supplementation markedly reduces cyst formation and restores tubule width toward normal levels in Drice mutant flies.

### 2.3. Cell number, shape and nucleus size gets restored upon BG supplementation in Drice mutants

Following the restoration of tubule morphology in *Drice* mutants upon BG supplementation, we quantified the different cell type numbers, examined their morphology and nuclear dimensions. The number of PCs in *Drice* mutants remained relatively unaltered after feeding with either 0.25% or 0.50% BG supplementation (Figure 3, B). In contrast, the number of SCs increased significantly in both BG-treated groups. SC numbers increased from 17.4±2.5 and 11.9±3.4 in the ATs and PTs respectively of Drice, NF group to 23.0 ±4.3 and 15.1±3.2 in 0.25% BG supplementation, and 28.6±4.5 and 14.5±2.3 in 0.50% BG supplemented *Drice* mutants, whereas, it was 30.7±3.0 and 20.3±2.3 in the ATs and PTs respectively in the wild type NF group (Figure 3, C; Supplementary Table S5).

**Figure 3:**
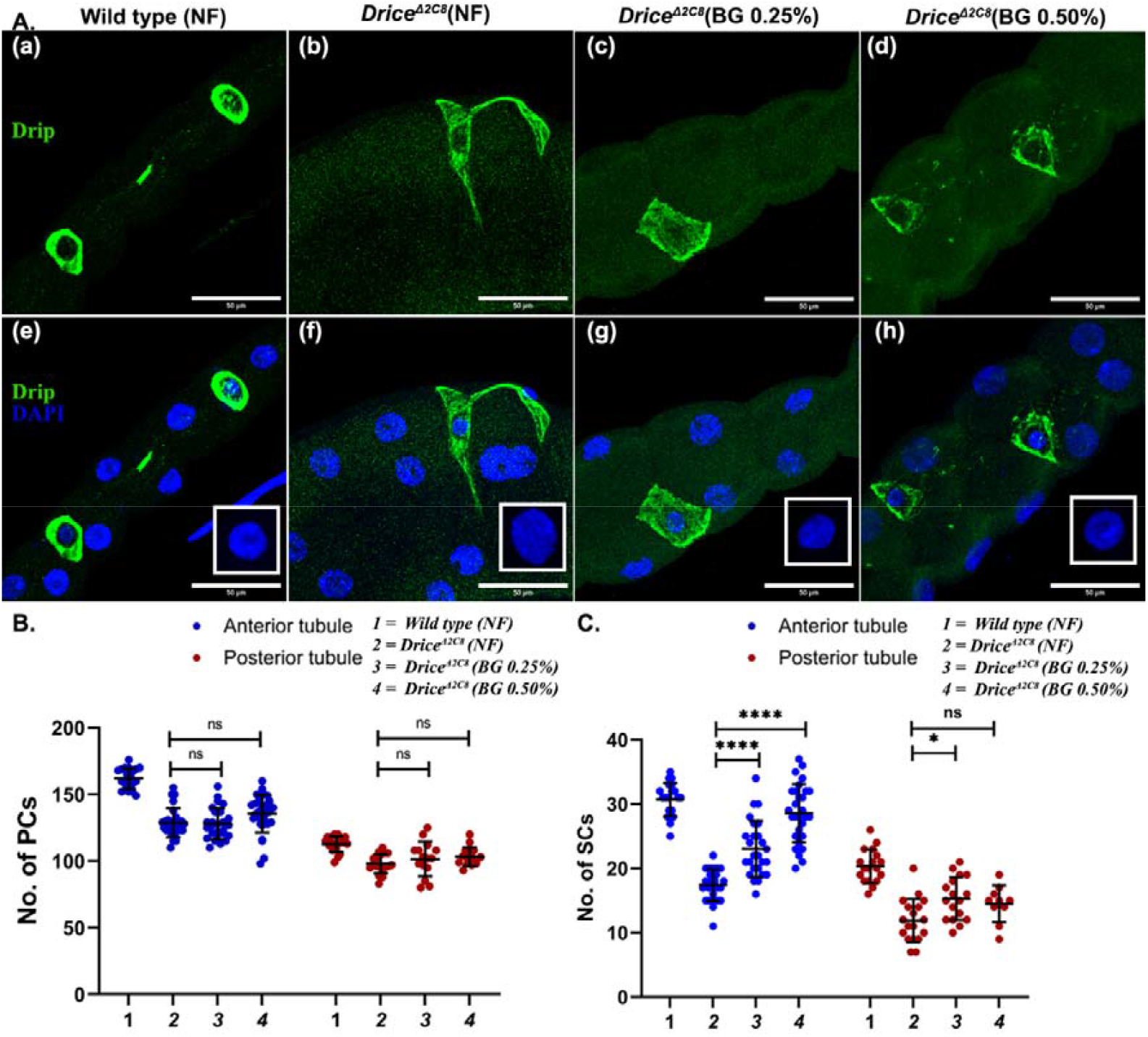
Restoration of SCs shape, size, and number in Drice mutants upon BG supplementation. (A) SC shape restoration. SCs in Drice mutants displayed irregular polygonal morphology (A-b, f) compared to the wild type (A-a, e). BG supplementation at 0.25% (A-c, g) and 0.50% (A-d, h) reduced aberrant protrusions and restored SCs to their typical cuboidal/rectangular shape, with more pronounced recovery at 0.25%. Insets (white boxes) show nuclear morphology, indicating reduced nuclear size upon BG supplementation. Scale bar = 50 µm. **(B) PCs number in MTs upon BG supplementation**. BG supplementation did not significantly alter PC number in either AT or PT: BG 0.25% (p = 0.9940, 0.7147 for AT and PT) and BG 0.50% (p = 0.1305, 0.4224 for AT and PT) compared to Drice mutants on NF (n ≥ 20 per group). **(C) BG supplementation restores the diminished SCs number in Drice deletion mutant**. A significant increase in the SCs number, with a stronger effect at BG 0.25% (p < 0.0001 for AT and PT) compared to BG 0.50% (p = 0.0364 for AT; p = 0.1138 for PT) (n ≥ 20 per group). Welch’s one way ANOVA test with Tukey’s post hoc test was done to determine the statistical significance. p-value < 0.05 is considered significant, with reference *p < 0.05, ** p <0.01, ***p < 0.001, and ****p < 0.0001. Bar graphs are showing Mean ± SEM value. n = 5 (technical replicate) * 3 independent biological replicates.

We next analyzed the shape of the SCs. Under normal conditions, SCs transition from cuboidal to star-shaped form during the larval-to-adult transition [17]. However, in *Drice* mutants, SCs prematurely acquire a star/polygonal shape during the larval stage itself [10]. BG feeding led to a partial restoration of SCs shape (Figure 3, A). In BG supplemented *Drice* mutants, the SCs that were star/polygonal-shaped in NF supplementation (Figure 3, A-b, f) reverted towards their typical cuboidal to rectangular morphology in both BG 0.25% (Figure 3, A-c, g) and BG 0.50% treatments (Figure 3, A-d, h) (refer to Supplementary Figure S8 for lower resolution).

Collectively, these findings indicate a substantial rescue of MT morphology in *Drice* mutants upon BG supplementation, primarily mediated through the restoration of SC number and shape.

### 2.4. BG supplementation show rescue in the cell polarity and cytoskeletal arrangement in the MTs of the Drice mutants

Previous studies have demonstrated that *Drice* mutants exhibit severe disorganization of F-actin and disruption of epithelial polarity due to aberrant activation of the Rho1 GTPase pathway [10] [11]. To assess whether BG supplementation could restore cytoskeletal and polarity defects, we analyzed F-actin organization, Dlg distribution, and Rho1 expression in *Drice* mutants following BG supplementation.

Confocal imaging revealed that F-actin in untreated *Drice* mutants was densely aggregated and highly disordered (Figure 4, A-b, f). Upon BG administration, F-actin organization was markedly restored, displaying a less compact and more filamentous arrangement comparable to wild-type tubules (Figure 4, A-a, e). Both 0.25% (Figure 4, A-c, g) and 0.50% (Figure 4, A-d, h) BG supplementation exhibited comparable efficacy in rescuing cytoskeletal organization. Further we also quantified F-actin organization by the Fourier-based directionality analysis revealed highly disorganized F-actin is getting restored upon BG supplementation (Figure 4, A, i).

**Figure 4:**
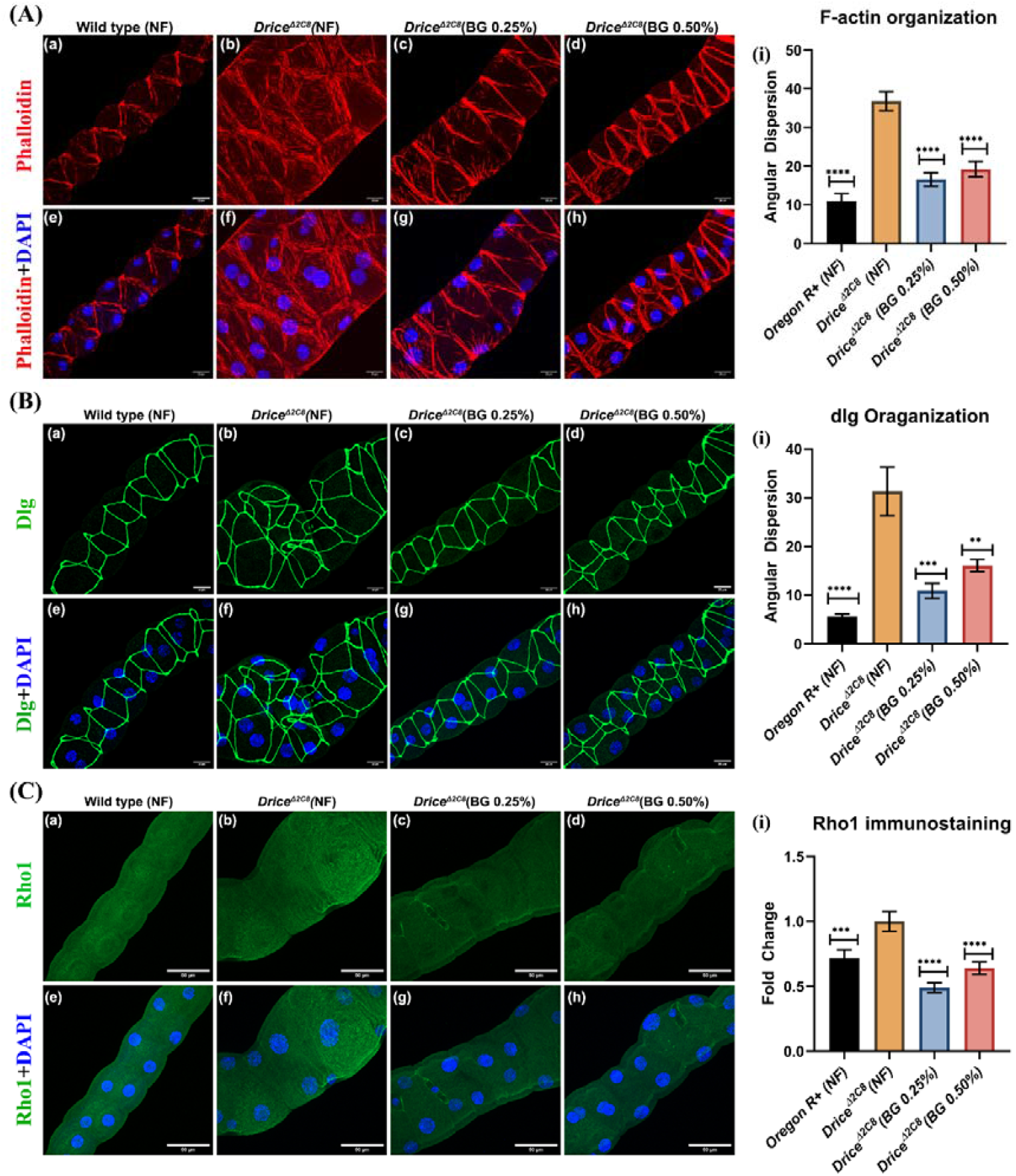
Effect of BG supplementation on actin organization, cell polarity, and Rho1 protein expression in *Drice* mutants. **(A) BG supplementation restored actin dynamics in Malpighian tubules (MTs) of *Drice* mutants**. Disorganized actin in mutants (A-b, f) became comparable to wild type (A-a, e) upon BG 0.25% (A-c, g) and BG 0.50% (A-d, h) supplementation. (A-i) Quantification of the actin organization (angular dispersion). F-actin is highly disorganized in Drice NF, group when compared to the wild type group. BG supplementation restores the F-actin organization in the MTs. Scale bar = 20 µm. **(B) BG supplementation restored apico-basal (A–P) cell polarity**, re-establishing the typical bilinear cellular arrangement in MTs at both BG 0.25% and BG 0.50% (B-c, g; B-d, h) compared to the disrupted polarity in *Drice* mutants (B-b, f). (A-i) Quantification of the cell polarity organization (angular dispersion). dlg is highly disorganized in Drice NF, group when compared to the wild type group. BG supplementation restores the cell-polarity in the MTs. Scale bar = 20 µm. **(C) BG supplementation downregulated Rho1 protein expression**, which was highly elevated in *Drice* mutants (C-b, f). Expression levels were reduced in both BG 0.25% (C-c, g) and BG 0.50% (C-d, h), becoming comparable to wild type (C-a, e) with (C-i) is showing the protein quantification. Scale bar = 50 µm.

Analysis of Dlg localization further supported the rescue of epithelial polarity. *Drice* mutants, NF showed profound polarity disruption, characterized by multilayered epithelial structures and loss of apical–basal organization (Figure 4, B-b, f). Both 0.25% (Figure 4, B-c, g) and 0.50% BG (Figure 4, B-d, h) re-established polarity, with Dlg confined to defined lateral membranes and cells arranged in a typical bilayer fashion, indicating effective restoration of epithelial integrity. Quantification of the polarity defects by the Fourier-based directionality analysis revealed very high polarity defects in Drice muatnts that are getting restored upon BG supplementation (Figure 4, B, i).

Given the established role of Rho GTPases in cytoskeletal regulation and podocyte foot process stability in mammalian nephrons [18]; and previous report of Rho1 mediated cytoskeletal disarray [10, 19]. We examined Rho1 protein expression in BG supplementation *Drice* mutants. Immunostaining revealed substantial downregulation of Rho1 levels in both 0.25% (Figure 4, C-c, g) and 0.50% BG supplementation (Figure 4, C-d, h) larvae relative to untreated controls (Figure 4, C-b & f; Figure 4, C-i).

Collectively, these findings demonstrate that BG supplementation restores F-actin organization and apico-basal polarity along with restoration of Rho1 GTPase protein expression in the MTs of the Drice mutants.

### 2.5. BG improves the physiology of the MTs in Drice mutant flies

Following the significant improvement in fly survival and Malpighian tubule (MT) morphology observed in *Drice* mutants upon *Brahmi Ghrita* (BG) supplementation, we next examined whether this morphological rescue was accompanied by restoration of normal tubule physiology. MTs in *Drice* mutants are known to exhibit compromised physiological function, as previously reported by our laboratory [10]. In insects, uric acid crystal formation and fluid secretion serve as reliable indicators of MT physiological status. Under normal conditions, uric acid crystals form transiently within the tubules but are subsequently cleared into the hindgut through peristaltic movements and eliminated along with fecal material [20].

Consistent with impaired excretory function, *Drice* mutants exhibited markedly elevated uric acid crystal deposition (Figure 5A-b, f, j) compared to wild-type controls (Figure 5A-a, e, i). Notably, BG supplementation significantly reduced crystal accumulation in the *Drice* mutant background at both 0.25% (Figure 5A-c, g, k) and 0.50% (Figure 5A-d, h, l) supplementation. These observations indicate a substantial improvement in MT physiological function following BG treatment.

**Figure 5:**
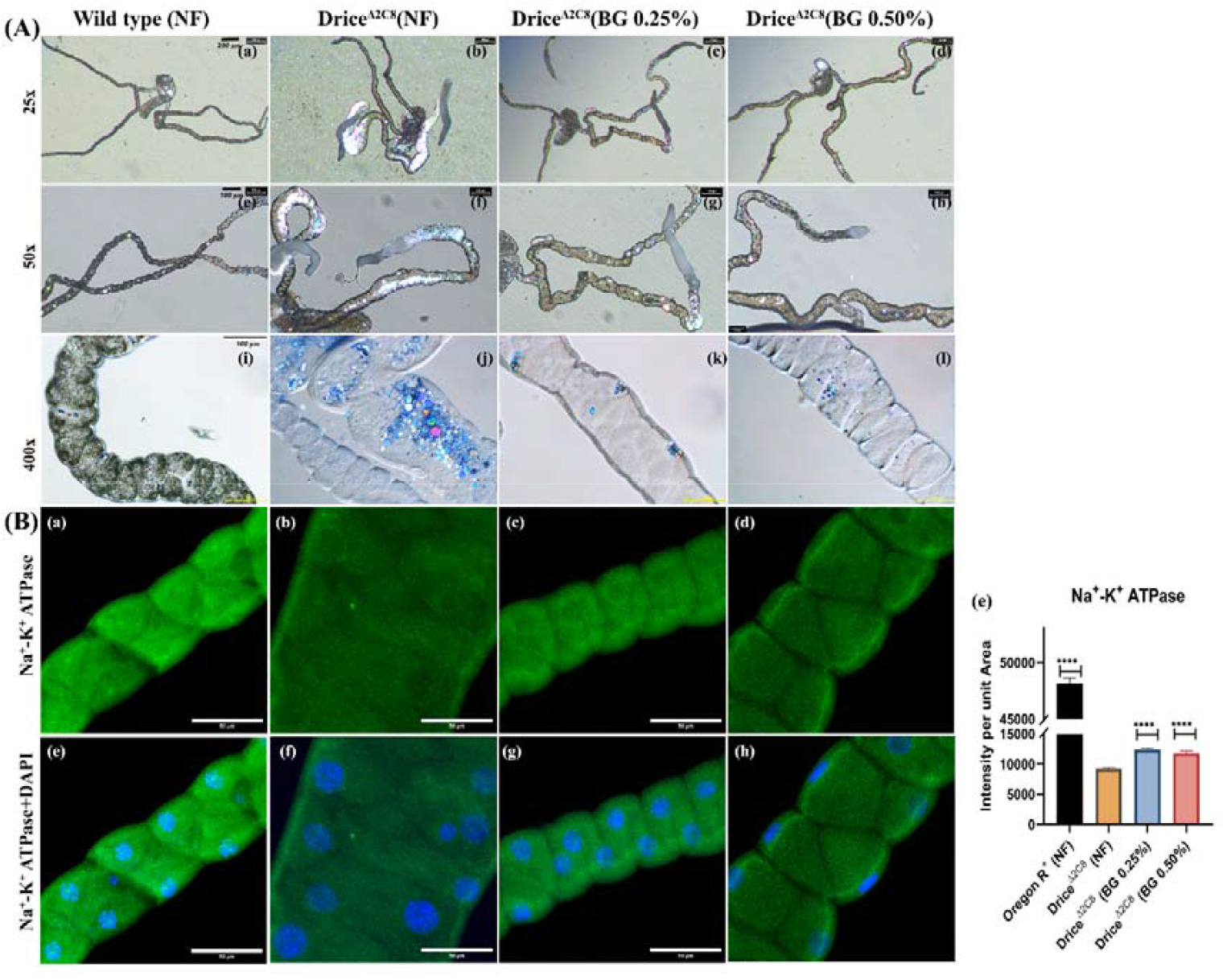
BG supplementation restores tubular function in *Drice* mutants. **(A) BG supplementation reduced uric acid crystal deposition in Malpighian tubules (MTs) of *Drice* mutants.** Mutants showed extensive uric acid accumulation (bright birefringent crystals under polarized light; b, f, j) compared to wild type (a, e, i). BG supplementation markedly reduced crystal deposition at both BG 0.25% (c, g, k) and BG 0.50% (d, h, l), indicating improved MT physiological function. Scale bars: 250 µm (a-h) and 100 µm (i-l). **(B) Immunostaining of Malpighian tubules showing Na□/K□-ATPase, and nuclear labeling across genotypes and dietary conditions**. Compared to wild-type tubules (a, e), Drice mutants maintained on normal food (NF) exhibit significantly reduced Gelsolin expression (b, f) and diminished Na□/K□-ATPase levels (i). Dietary supplementation with Brahmi Ghrita (BG) at 0.25% (c, g) and 0.50% (d, h) markedly restores Gelsolin expression and enhances Na□/K□-ATPase levels in Drice mutant tubules relative to NF group. DAPI marks nuclei. Welch’s one way ANOVA test with Tukey’s post hoc test was done to determine the statistical significance. p-value < 0.05 is considered significant, with reference *p < 0.05, ** p <0.01, ***p < 0.001, and ****p < 0.0001. Bar graphs are showing Mean ± SEM value. n = 5 (technical replicate) * 3 independent biological replicates.

To further substantiate these findings, we next examined the status of the Na□/K□-ATPase, a key ion transporter that provides the electrochemical driving force for epithelial transport and maintains the ionic gradients required for fluid secretion and filtration [21-23]. Immunostaining analyses revealed that Na□/K□-ATPase levels were markedly reduced in *Drice* mutants (Figure 6, B-a & e) compared to wild-type tubules (Figure 6, B-b & f). Importantly, BG supplementation significantly enhanced Na□/K□-ATPase expression in *Drice* mutants at both 0.25% (Figure 6, B-c & g) and 0.50% (Figure 6, B-d & h) concentrations (Figure 6, B-i). These results indicate that the restoration of tubular fluid secretion observed in BG-treated *Drice* mutants is likely mediated, at least in part, through the recovery of Na□/K□-ATPase–dependent ion transport. These results further support the conclusion that BG supplementation effectively rescues MT physiological function in the *Drice* mutant background.

**Figure 6.**
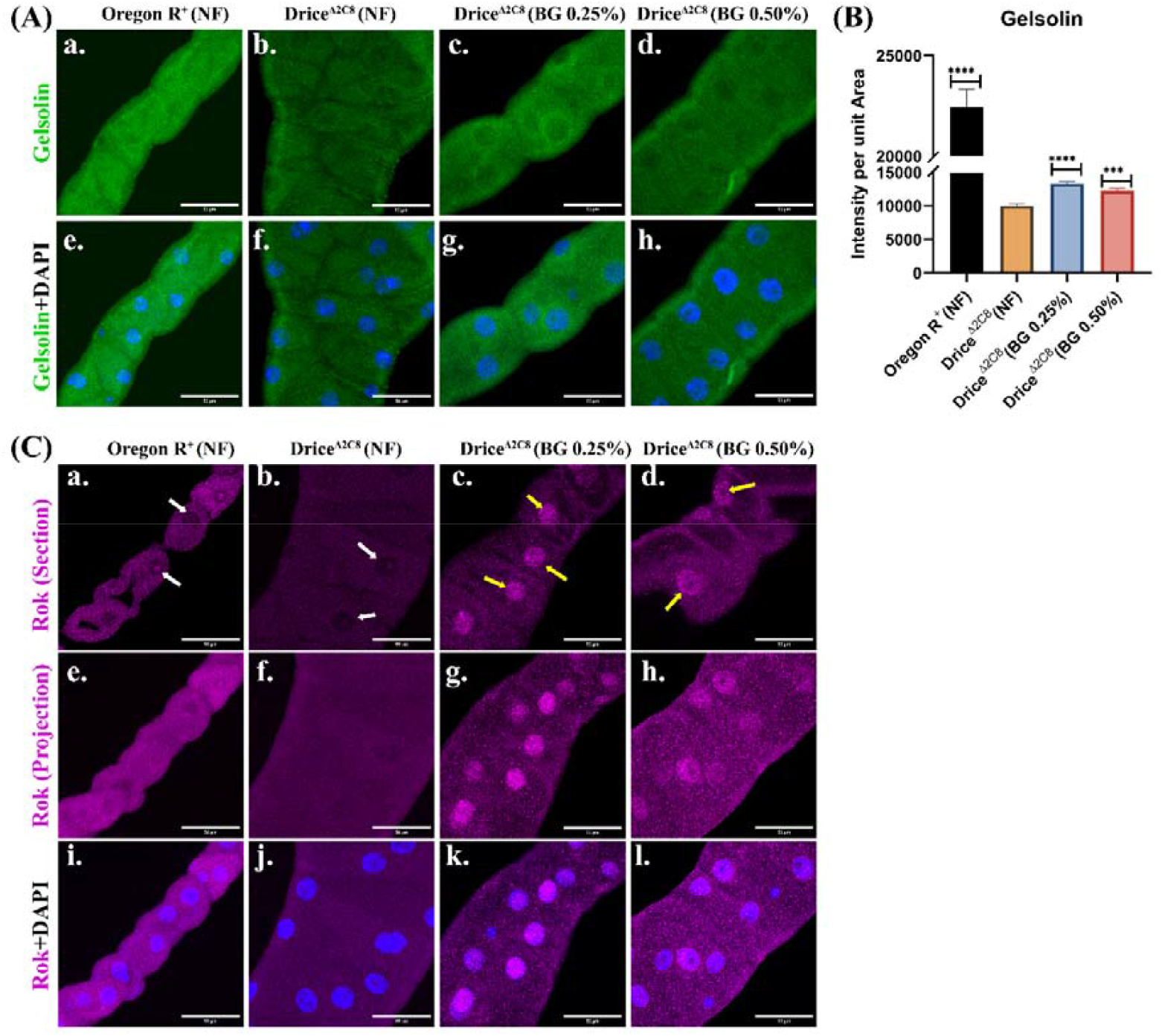
BG supplementation improves Gelsolin levels and promotes the Rok nuclearization in the MTs of the Drice deletion mutants. **(A) Immunostaining of Malpighian tubules showing expression of Gelsolin protein.** Compared to wild-type tubules (a, e), Drice mutants maintained on normal food (NF) exhibit significantly reduced Gelsolin expression (b, f). Dietary supplementation with Brahmi Ghrita (BG) at 0.25% (c, g) and 0.50% (d, h) markedly restores Gelsolin expression levels in Drice mutant tubules relative to NF group. DAPI marks nuclei. **(B) Quantification** of fluorescence intensity confirms significant recovery of Gelsolin upon BG supplementation. Welch’s one way ANOVA test with Tukey’s post hoc test was done to determine the statistical significance. p-value < 0.05 is considered significant, with reference *p < 0.05, ** p <0.01, ***p < 0.001, and ****p < 0.0001. Bar graphs are showing Mean ± SEM value. n = 5 (technical replicate) * 3 independent biological replicates. **(C) BG supplementation promotes Rok nuclearization in the MTs of the Drice deletion mutants**. Drice mutant, NG group (b, f & j) are showing reduced Rok levels compared to the wild type, NF group (a, e & i) as reported earlier. Both BG 0.25% (c, g & k) and BG 0.50% (d, h & l) supplementation promotes the translocation of the Rok into the nucleus of the MTs in Drice deletion mutants. (a-d are showing the section images, -l are showing the projection image).

### 2.6: BG supplementation improves the expression of actin severing protein and transmembrane ion transport protein in the MTs of Drice deletion mutants

Following the previously reported by our laboratory, the dense accumulation of F-actin observed in *Drice* mutants results from the downregulation of Gelsolin, an actin-severing and capping protein that prevents excessive F-actin buildup within the cellular environment [19]. Consistent with earlier findings, Gelsolin protein levels were significantly reduced in *Drice* mutants maintained on NF (Figure 6, A-b, f) compared to the wild-type genotype (Figure 6, A-a, e) [19]. Notably, BG supplementation led to a marked restoration of Gelsolin expression in *Drice* mutants at both 0.25% (Figure 6, A-c, g) and 0.50% (Figure 6, A-d, h) concentrations relative to NF group (Figure 6, A-b, f) (Figure 6B). These findings suggest that improved actin organization and restoration of cell polarity in BG-treated *Drice* mutants may be associated with the recovery of Gelsolin protein levels.

The disruption of actin dynamics in the absence of caspase-3 activity is associated with the failure of Rok activation, despite the high abundance of Rho1 protein [19]. Based on this observation, we next examined G-actin and Rok levels in the Drice deletion mutant upon BG supplementation. Rok exhibited a marked change in subcellular distribution upon BG treatment. In wild-type, NF (Figure 7, C-a, e & i) and the Drice NF group (Figure 7, C-b, f & j), Rok localization was predominantly cytoplasmic. However, BG supplementation at both 0.25% (Figure 7, C-c, g & k) and 0.50% (Figure 7, C-d, h & l) concentrations resulted in prominent nuclear enrichment of Rok in Drice mutant tubules. However, due to technical limitations, the downstream transcriptional targets of Rok could not be explored in the present study. Nonetheless, we hypothesize that Rok nuclearization, together with Gelsolin and Na□/K□-ATPase, contributes to the rescue of morphological and physiological defects resembling the PKD phenotype in Drice deletion mutants.

## 3. Discussion

Polycystic kidney disease (PKD) is a genetically inherited disorder characterized by progressive cyst formation driven by aberrant epithelial proliferation, dysregulated fluid secretion, and disruption of tissue architecture, ultimately leading to end-stage renal disease (ESRD) [1, 24]. A striking hallmark of PKD is the excessive radial expansion of epithelial tubules, wherein a normal renal tubule of ∼40 µm diameter can enlarge to several centimeters in advanced cystic kidneys [25]. Several fundamental aspects of renal tubule biology—including epithelial polarity, cytoskeletal organization, and ion and water transport—are evolutionarily conserved between mammalian nephrons and the Malpighian tubules (MTs) of Drosophila melanogaster [26-28]. This conservation underscores the utility of Drosophila MTs as a robust in vivo system for dissecting the cellular mechanisms underlying renal cystogenesis and for evaluating potential therapeutic interventions for PKD.

Our laboratory previously demonstrated that loss of caspase-3/Drice in Drosophila results in a pronounced PKD-like phenotype in MTs, characterized by cystic dilation, disorganized cytoskeleton, loss of epithelial polarity, and impaired excretory physiology [10, 19]. In the present study, we show that the polyherbal Ayurvedic Rasayana formulation Brahmi Ghrita (BG) significantly ameliorates these PKD-like defects in Drice homozygous mutants, leading to improved organismal survival and restoration of both tubular architecture and physiological function.

BG is traditionally recognized in Ayurveda as a Medhya Rasayana with established neuroprotective and anticonvulsant properties [13-15], and has recently been reported to exert nephroprotective effects in mammalian models of drug-induced renal toxicity [16]. Given its polyherbal composition, the present study does not seek to identify a single bioactive component responsible for the observed effects. Rather, our data support a model in which BG exerts a functional convergence on cytoskeletal remodeling and epithelial transport machinery. In Drice mutant MTs—typically marked by extensive cystic dilation, disrupted cellular organization, and defective excretion—BG supplementation resulted in a marked reduction in cyst formation and normalization of tubule diameter, while tubule length remained unchanged. This selective correction of tubule width is consistent with restoration of epithelial organization rather than altered developmental growth.

At the cellular level, the rescue of MT morphology was closely associated with restoration of stellate cell (SC) number and morphology. SCs play a central role in regulating water flux, chloride conductance, and rapid physiological adaptation of MTs, whereas principal cells (PCs) primarily mediate basal ion transport and epithelial organization [20, 21, 29]. BG supplementation significantly increased SC number and partially restored their aberrant polygonal/star-shaped morphology toward a cuboidal–rectangular configuration characteristic of larval stages. Importantly, PC numbers remained unchanged, indicating that BG does not induce global cellular hyperplasia. Instead, improved MT physiology is likely driven by restoration of SC abundance and morphology, together with recovery of PC-associated transport machinery.

Consistent with this interpretation, BG supplementation markedly enhanced Na□/K□-ATPase expression, a key driver of epithelial ion gradients and fluid secretion. Restoration of this transporter provides a mechanistic link between improved epithelial architecture and the observed recovery of tubule function, including reduced uric acid crystal accumulation. Together, these findings indicate that BG-mediated rescue involves coordinated restoration of both cellular composition and ion transport capacity, culminating in functional recovery of the renal analog.

Disruption of cytoskeletal organization and epithelial polarity is a central driver of cystic dilation in PKD. MT remodeling during development is governed by tightly regulated convergent extension processes that depend on apico–basal and proximal–distal polarity cues [30]. In Drice mutants, aberrant accumulation and disorganization of F-actin, together with mislocalization of polarity determinants such as Disc large (dlg), lead to loss of epithelial integrity and abnormal tubule architecture [10]. BG supplementation effectively restored F-actin organization and re-established lateral Dlg localization, indicating recovery of epithelial polarity. Restoration of polarity likely limits aberrant cell clustering, promotes re-establishment of a bilinear epithelial arrangement, and contributes directly to normalization of tubule diameter.

At the molecular level, Drice loss is known to dysregulate Rho1 GTPase signaling, resulting in cytoskeletal instability and polarity defects [10, 19]. BG supplementation significantly reduced elevated Rho1 protein levels in Drice mutants, consistent with normalization of actin dynamics. Notably, BG selectively restored expression of Gelsolin, an actin-severing and capping protein whose downregulation in Drice mutants drives pathological F-actin accumulation. Recovery of Gelsolin provides a direct mechanistic explanation for the observed correction of actin organization and epithelial polarity. Interestingly, although total Rok protein levels show a pronounced shift in subcellular localization following BG supplementation. In untreated Drice mutants, Rok was predominantly cytoplasmic, consistent with its established role in actomyosin regulation. Upon BG supplementation, Rok exhibited significant nuclear enrichment, suggesting altered compartmentalization of Rok signaling in the mutant epithelium.

Rok family kinases are classically recognized for cytoplasmic regulation of contractility and cytoskeletal dynamics. However, nuclear localization of Rok has been reported under specific developmental and stress-responsive contexts, including roles in transcriptional modulation and cell-cycle control [31-34]. Whether the nuclear enrichment of Rok observed here reflects transcription-associated activity, regulatory feedback to Rho signaling, or altered cytoskeletal equilibrium remains unresolved. Importantly, the present study does not establish a causal hierarchy between Rok redistribution, Gelsolin recovery, Rho1 normalization, or Na□/K□-ATPase restoration. Rather, these events occur concomitantly in the BG-treated Drice mutant background. Future genetic epistasis experiments, activity assays, and transcriptional analyses will be required to determine pathway dependency and functional significance of nuclear Rok.

Collectively, our findings support a working model in which BG supplementation coincides with normalization of Rho1 protein levels, restoration of Gelsolin-mediated actin remodeling, and redistribution of Rok to the nucleus, together facilitating recovery of epithelial organization and transport function. This multifaceted rescue results in reduced cyst formation, improved tubule physiology, and enhanced organismal survival. Nevertheless, the present work establishes Brahmi Ghrita as a potent modulator of epithelial integrity and renal analog function, and highlights the utility of Drosophila MTs as a powerful platform for mechanistic evaluation of traditional formulations in PKD-relevant contexts.

## 4. Material and methods

### 4.1.1. Drosophila stock husbandry

*Drosophila melanogaster* stocks were maintained on a standard fly medium consisting of 4.6% cornmeal, 4.5% sucrose, 1.6% yeast extract, and 0.7% agar, supplemented with 0.3% propionic acid and 0.3% methyl p-hydroxybenzoate as preservatives. All fly stocks were maintained at room temperature, while experimental crosses and assays were conducted at 24 °C under a 12 h light/12 h dark photoperiod. The following Drosophila stocks were used in this study: w; +/+; +/+ (wild-type strain), w^111^□; +/+; +/+, and w^111^□; +/+; DriceΔ2C8/TM6B.

### 4.2. Composition of BG

BG was procured in ready to use form from Arya Vaidya-Sala Kottakkal, Kerala, India (HPTLC data is attached; Supplementary Figure S8) with the composition per 100 g in the supplementary file.

### 4.3. Supplementation food preparation

Standard fly food was prepared as described above and allowed to cool to approximately 45 °C before supplementation. *Brahmi Ghrita* (BG), an Ayurvedic *Rasayana* formulation, was then added to the medium at final concentrations of 0.10%, 0.25%, 0.50%, and 1.0% (w/v). These concentrations were selected based on previously reported dietary supplementation ranges used for herbal and lipid-based formulations in *Drosophila*, as well as to assess potential dose-dependent effects while avoiding adverse impacts on food palatability, larval development, and adult viability.

### 4.4. Longevity Assay

First-instar larvae collected within one hour of hatching were transferred to food containing the indicated concentrations of supplementation or to normal food (control). To minimize experimental variability, control and BG-supplemented food were prepared from the same batch and larvae from the same cohort were used across all experimental groups. For each condition, approximately *n* = 25–30 flies were maintained per vial, with a minimum of three independent biological replicates per genotype and treatment. Adult flies were transferred to fresh vials every alternate day without the use of anesthesia, and the number of surviving flies was recorded at each transfer until all individuals had died or until 60 days of age.

### 4.5. Fecundity Assay

Male and female flies were separated within one hour of eclosion under brief ether anesthesia and maintained separately in individual vials for three days. Mating was then established in a 2 × 2 factorial design, comprising two males and two females per vial, for both control and BG-supplemented feeding groups. For each genotype and dietary condition, a minimum of 8–10 independent mating vials were set up as biological replicates. Flies were transferred to fresh vials daily, and the number of eggs laid was counted and recorded every day for up to 45 days.

### 4.6. MT length measurement and cell counting

Malpighian tubules (MTs) were dissected in chilled 1× phosphate-buffered saline (PBS) under a stereo-binocular microscope. Bright-field images of the dissected MTs were captured using a Nikon SMZ800N stereo-binocular microscope at 15× magnification. Tubule length was subsequently measured using ImageJ software. For cell counting, MTs were dissected, fixed, and stained with DAPI, a nuclear-specific fluorescent dye. Stained preparations were examined using a Nikon ECLIPSE E800 fluorescence microscope. Principal cells (PCs) and stellate cells (SCs) were distinguished based on their characteristic nuclear size and morphology and quantified using 20× (200× total magnification) and/or 40× (400× total magnification) objectives.

### 4.7. MTs dissection and Immunostaining

Malpighian tubules (MTs) were dissected from wandering third-instar larvae in 1× phosphate-buffered saline (PBS). Immediately following dissection, tissues were fixed in 4% paraformaldehyde (PFA) for 45 min at room temperature. After fixation, samples were washed with 0.5% PBST (0.5% Triton X-100 in 1× PBS) and subsequently incubated in blocking solution for 2 h at room temperature. The blocking solution consisted of 0.1% Triton X-100, 0.1% bovine serum albumin (BSA), 10% fetal bovine serum (FBS), 0.1% deoxycholate, and 0.02% thiomersal.

Following blocking, tissues were incubated overnight with primary antibodies at 4 °C. After primary antibody incubation, samples were washed three times with PBST and re-blocked for 2 h at room temperature before incubation with the appropriate secondary antibodies for 2 h under identical conditions. Tissues were then washed again with 0.5% PBST and counterstained with DAPI (4′,6-diamidino-2-phenylindole dihydrochloride; Thermo Fisher Scientific, Cat. #D1306) at a final concentration of 1 μg/mL to visualize nuclei.

After three final washes, samples were mounted on glass slides using DABCO mounting medium, covered with glass coverslips, and sealed with clear nail polish. Slides were stored at 4 °C for short-term use and at −20 °C for long-term storage. All primary and secondary antibodies were diluted in blocking buffer.

The following primary antibodies were used:

anti-dlg antibody raised in mouse (1:100, 4f-3, DSHB),
anti-Rho1 antibody raised in rabbit (1:1500, 10749-1-AP, Protein tech),
anti-Rho1 antibody raised in mouse (1:1000, SC-0418, Santa Cruz),
anti-Drip antibody raised in rabbit (1:1000, Kind gift from Dr. J. A. T. Dow)
anti-Rok antibody raised in rabbit (1:1000, Gly1114, CST),
anti-Gelsolin antibody raised in rabbit (1:1000, D9W8Y, CST),
The secondary antibodies used for the immunohistochemistry are as follows:
donkey anti-mouse AF 555 (A31570),
goat anti-mouse AF 647 (A21050),
donkey anti-rabbit AF 555 (A31572) and
goat anti-rabbit AF 647 (A32733) from Invitrogen.

### 4.8. Phalloidin staining

Malpighian tubules (MTs) were dissected from 118–120 h old larvae at the inverted spiracle stage in chilled 1× phosphate-buffered saline (PBS) and immediately fixed in 4% paraformaldehyde (PFA) for 45 min. Following fixation, tissues were washed three times in 0.5% PBST (0.5% Triton X-100 in 1× PBS) at room temperature and incubated with phalloidin stain for 2 h at room temperature. After staining, samples were washed again in 0.5% PBST and counterstained with DAPI for 20 min at room temperature. Tissues were subsequently washed three times in 0.5% PBST, mounted in DABCO mounting medium on glass slides, and imaged using a fluorescence microscope.

### 4.9. DNaseI staining

Malpighian tubules (MTs) were dissected from 118–120 h old larvae at the inverted spiracle stage in chilled 1× phosphate-buffered saline (PBS) and immediately fixed in 4% paraformaldehyde (PFA) for 45 min. Following fixation, tissues were washed three times in 0.5% PBST (0.5% Triton X-100 in 1× PBS) at room temperature and incubated with DNase I stain for 20 min at room temperature. After staining, samples were washed again in 0.5% PBST and counterstained with DAPI for 20 min at room temperature. Tissues were subsequently washed three times in 0.5% PBST, mounted in DABCO mounting medium on glass slides, and visualized using a fluorescence microscope.

### 4.10. Scanning electron microscopy

Malpighian tubules (MTs) were dissected from 118– 120 h old larvae at the inverted spiracle stage in chilled 1× phosphate-buffered saline (PBS) and immediately fixed in a 1:1 mixture of 4% paraformaldehyde and glutaraldehyde for 30 min at room temperature. Following fixation, tissues were washed three times in 1× PBS (5 min each) at room temperature. Samples were then dehydrated through a graded ethanol series consisting of 30%, 50%, 70%, 90%, and 100% ethanol, followed by absolute ethanol, with two changes at each step for 5 min at room temperature. After dehydration, tissues were air-dried in a desiccator for 1–2 days until complete removal of moisture. Dried samples were mounted on aluminum stubs using conductive carbon tape, sputter-coated with gold nanoparticles, and examined using a Gemini-SEM 560 scanning electron microscope.

### 4.11. Polarizing microscopy

Malpighian tubules (MTs) were dissected in chilled 1× phosphate-buffered saline (PBS) and immediately mounted on glass slides under low-temperature conditions. The mounted tissues were examined without delay using a polarized light microscope to visualize the birefringent (glittering) uric acid crystals within the MTs.

### 4.12. Microscopy and image processing

All samples were imaged using a Zeiss LSM 900 confocal microscope with a 20× objective (zoom 1.0) or a 40× objective (zoom 0.8). Optical sections were acquired at a z-interval of 2.0 µm for all samples, unless otherwise specified in the figure legends. Image acquisition parameters were kept constant across experimental groups within each experiment. All images were processed and analyzed using ImageJ software (NIH, USA; https://imagej.nih.gov/ij), and figure panels were assembled using Adobe Photoshop (Version 22.4.2). Images of Malpighian tubules are presented as maximum-intensity projections of the complete z-stacks, enabling accurate visualization of the tubule lumen and epithelial architecture. Given the hollow nature of Malpighian tubules, single optical sections were not used for general morphological analysis unless specifically indicated.

### 4.13. Statistical analysis

All experiments were performed with a minimum of three independent biological replicates unless otherwise stated. For each biological replicate, at least 10 Malpighian tubules (MTs) or tissues were analyzed per experimental condition. Data are presented as mean ± standard error of the mean (SEM).

Statistical analyses were performed using GraphPad Prism (version 8.4.2). For comparisons between two groups, unpaired two-tailed Student’s *t*-tests were used. When multiple pairwise comparisons were performed, Bonferroni correction was applied where appropriate. For comparisons involving more than two groups, one-way analysis of variance (ANOVA) followed by Tukey’s multiple-comparison post hoc test was employed. In cases where variance was unequal among groups, Welch’s one-way ANOVA was used. Two-way ANOVA followed by Tukey’s post hoc test was applied for experiments involving two independent variables.

For survival analyses (longevity assays), Kaplan–Meier survival curves were generated and statistical significance was determined using the log-rank (Mantel–Cox) test.

p-value ≤ 0.05 was taken as significant difference and degree was significance was as follow: p-values ≤0.05, ≤0.01, ≤0.001, and ≤0.0001 are signifying*, **, ***, and ****, respectively; with p-value > 0.05 representing non-significant statistical difference.

## Abbreviations

AF: Alexa Fluor
ATs: Anterior Tubules
BG: Brahmi Ghrita
BSA: Bovine Serum Albumin
DABCO: 1,4-Diazabicyclo[2.2.2]octane
DAPI: 4′,6-Diamidino-2-phenylindole dihydrochloride
DIC: Differential Interference Contrast
Dlg: Discs large
Drice: Drosophila interleukin-1β–converting enzyme (Caspase-3)
EMT: Epithelial-to-Mesenchymal Transition
ESRD: End-Stage Renal Disease
F-actin: Filamentous actin
FBS: Fetal Bovine Serum
G-actin: Globular actin
GFP: Green Fluorescent Protein
HPTLC: High-Performance Thin-Layer Chromatography
LC50: Lethal Concentration 50%
MTs: Malpighian Tubules
NF: Normal Food
no.: Number
PBS: Phosphate-Buffered Saline
PBST: Phosphate-Buffered Saline with Triton X-100
PCs: Principal Cells
PKD: Polycystic Kidney Disease
PMSF: Phenylmethylsulfonyl Fluoride
PTs: Posterior Tubules
Rho1: Ras homolog family member 1
Rok: Rho-associated kinase
RT: Room Temperature
SCs: Stellate Cells
SEM: Scanning Electron Microscopy / Standard Error of the Mean (use contextually)
SDS: Sodium Dodecyl Sulfate
TM6B: Third Multiple balancer chromosome (Drosophila)

## 6. Acknowledgements

We acknowledge the Department of Zoology, Institute of Science, Banaras Hindu University, Varanasi, for providing basic facility and infrastructure and Central Equipment Facility (CEF). We thank Dr. J. A. T. Dow, Institute of Biomedical Science, University of Glasgow for anti-Drip antibody, and Dr. M. Miura, Department of Genetics, University of Tokyo; for DriceΔ2c8 stocks. We acknowledge the Interdisciplinary School of Life Sciences & central discovery center (CDC) SATHI BHU, BHU, for Confocal microscopy facility. We acknowledge central discovery center (CDC) SATHI BHU, for IP-MS facility. We thank Dr. Bama Charan Mandal, Department of Zoology, Banaras Hindu University, Varanasi for Confocal microscopy facility. We thank Dr. Ajit Kumar Sahoo, Department of Geology, Banaras Hindu University for providing the polarizing microscope facility. Credit is also attributed to BioRender, for Graphical abstract and Work summary creation.

## 7. Author’s Contribution: Mr. Saurabh Chand Sagar

Conceptualization, Validation, Visualization, Writing – original draft, Writing – review and editing, Investigation, Methodology, Data curation, Conceptualization, Formal analysis, Analysis, Interpretation. **Dr. Madhu G. Tapadia**: Writing – review & editing, Validation, Supervision, Conceptualization, Funding acquisition, Project administration, Resources, Supervision, Interpretation.

