## Supplementary File for "Brahmi Ghrita Exerts Nephroprotective Effects by Restoring Cytoskeletal Integrity and Ion Transport in *Drosophila* Model of Polycystic Kidney Disease"

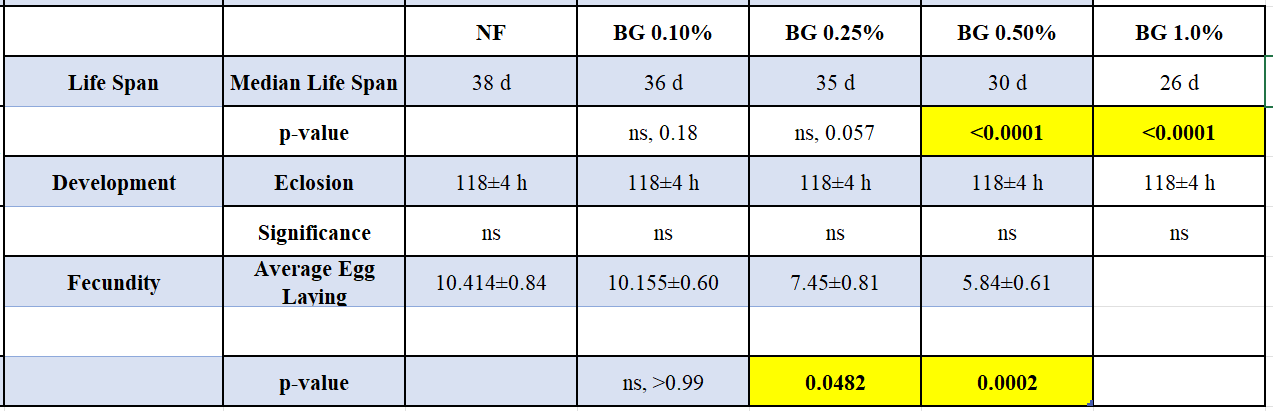

**Supplementary Table S1: Effect of Brahmi Ghrita (BG) supplementation on lifespan, development, and fecundity in wild-type *Drosophila melanogaster*.** Dietary supplementation with Brahmi Ghrita was well tolerated at lower concentrations, with BG 0.10% and 0.25% showing no significant effects on median lifespan or developmental timing, as assessed by eclosion rate. In contrast, higher concentrations (0.50% and 1.0%) produced a significant reduction in lifespan, indicating dose-dependent effects at elevated levels. Fecundity remained unaffected at BG 0.10%, while BG 0.25% resulted in a mild but statistically significant reduction, and higher concentrations caused a pronounced decrease in egg-laying capacity. These findings demonstrate that BG exhibits a defined tolerability window in *Drosophila*, supporting the use of BG 0.25% as an effective and physiologically acceptable dose for mechanistic and therapeutic evaluation. n = 800 per group for longevity, 500 per group for developmental assay, 20 pairs per group for fecundity assay. Statistical analysis for longevity, Kaplan–Meier survival analysis followed by the log-rank (Mantel–Cox) test, and Welch’s one-way ANOVA followed by Dunnett’s multiple-comparison test for development and fecundity was performed.

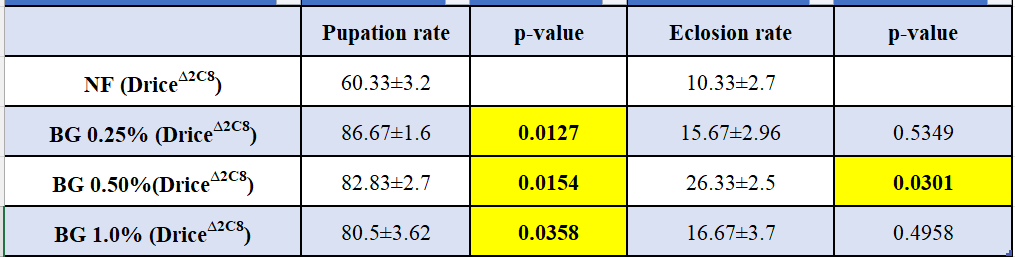

**Supplementary Table S2: Effect of Brahmi Ghrita (BG) supplementation on pupation and eclosion efficiency in *Drice*Δ2C8 mutant *Drosophila*.** Pupation and eclosion rates were quantified in *Drice*^Δ2C8^ mutants reared on normal food (NF) or BG-supplemented diets. Data are presented as mean ± SEM. Statistical comparisons were performed using Welch’s one-way ANOVA followed by Dunnett’s multiple-comparison test, with NF-fed mutants serving as the reference group. BG supplementation significantly improved pupation efficiency at all tested concentrations, with the strongest effect observed at BG 0.25% (*p* = 0.0127) and BG 0.50% (*p* = 0.0154). Eclosion efficiency showed a dose-dependent response, with a significant increase observed at BG 0.50% (*p* = 0.0301), whereas BG 0.25% and BG 1.0% did not differ significantly from NF controls. These findings indicate that BG enhances developmental progression and viability in *Drice* mutants within a defined dose window.

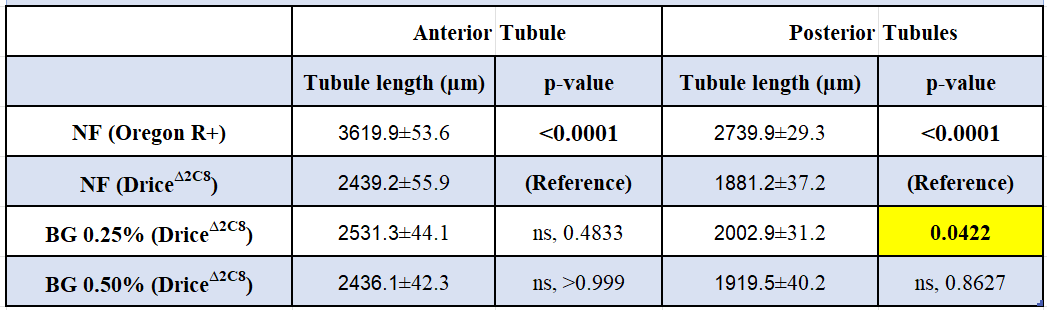

**Supplementary Table S3: Quantitative analysis of Malpighian tubule length in control and Drice mutant backgrounds with BG treatment.** Quantification of anterior and posterior Malpighian tubule (MT) lengths in control flies (NF, Oregon R+) and in *Drice*^Δ2C8^ mutants under normal feeding (NF) conditions or following BG treatment (0.25% and 0.50%). Data are presented as mean ± SEM (µm). For both anterior and posterior tubules, NF (Oregon R+) values are significantly longer than *Drice*^Δ2C8^ mutants (p < 0.0001). BG treatment (0.25%) in the *Drice*^Δ2C8^ background results in a modest but significant increase in posterior tubule length (p = 0.0422), while no significant change is observed in anterior tubule length (p = 0.4833). Treatment with 0.50% BG does not significantly alter tubule length in either anterior or posterior segments (ns, p > 0.05). Statistical comparisons were performed using one-way ANOVA with post hoc analysis, with *Drice*^Δ2C8^ NF serving as the reference group for treated conditions.

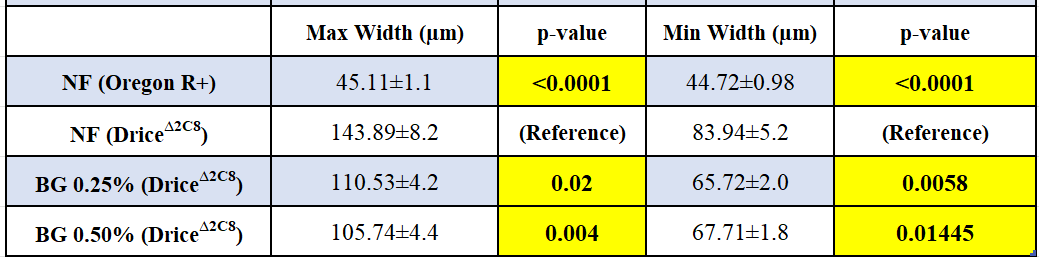

**Supplementary Table S4: Quantitative analysis of Malpighian tubule width in control and *Drice* mutant backgrounds with BG treatment.** Maximum and minimum widths (µm) of Malpighian tubules (MTs) were measured in control flies (NF, Oregon R+) and in *Drice*^Δ2C8^ mutants under normal feeding (NF) conditions or following BG supplementation (0.25% and 0.50%). Data are presented as mean ± SEM. Compared with NF (Oregon R+) controls, *Drice*^Δ2C8^ mutants exhibit a marked increase in both maximum and minimum tubule width (p < 0.0001 for both). Treatment of *Drice*^Δ2C8^ mutants with BG results in a significant reduction in tubule width. At BG 0.25%, both maximum width (p = 0.02) and minimum width (p = 0.0058) are significantly decreased relative to untreated *Drice*^Δ2C8^ mutants. Similarly, BG 0.50% treatment significantly reduces maximum width (p = 0.004) and minimum width (p = 0.01445). Statistical comparisons were performed using one-way ANOVA with post hoc analysis, with untreated *Drice*^Δ2C8^ (NF) serving as the reference group for treated conditions.

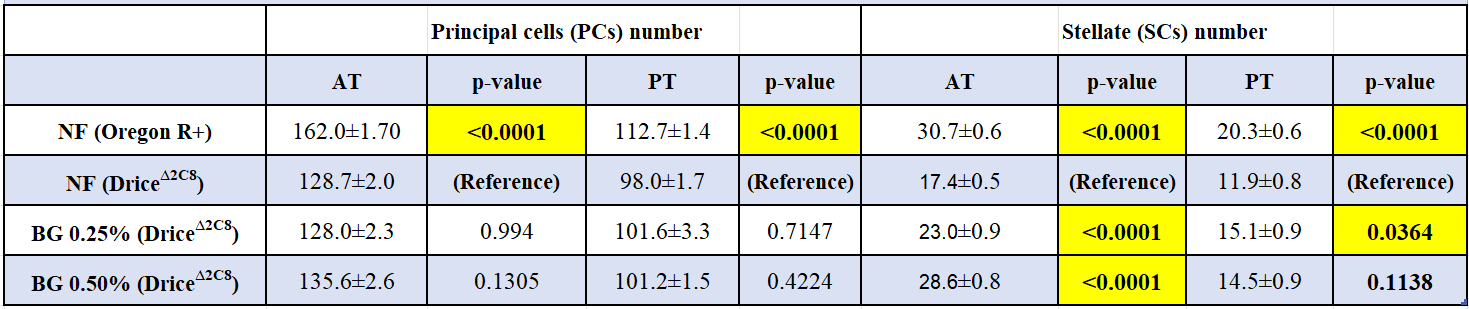

**Supplementary Table S5: Quantification of principal and stellate cell numbers in anterior and posterior Malpighian tubules under control and *Drice* mutant conditions with BG treatment.** Quantification of principal cells (PCs) and stellate cells (SCs) in anterior (AT) and posterior (PT) Malpighian tubules (MTs) from control flies (NF, Oregon R+) and mutants under normal feeding (NF) or BG treatment (0.25% and 0.50%). Data are presented as mean ± SEM. Compared with NF (Oregon R+) controls, *Drice*^Δ2C8^ mutants exhibit a significant reduction in both PC and SC numbers in anterior and posterior tubules (p < 0.0001 for all comparisons). BG treatment does not significantly alter principal cell numbers in either anterior or posterior tubules relative to untreated *Drice*^Δ2C8^ mutants (p > 0.05). In contrast, BG treatment significantly increases stellate cell numbers in anterior tubules at both 0.25% and 0.50% concentrations (p < 0.0001), while a modest but significant increase is observed in posterior tubules at 0.25% BG (p = 0.0364), but not at 0.50% BG (p = 0.1138). Statistical comparisons were performed using one-way ANOVA with post hoc analysis, with untreated *Drice*^Δ2C8^ (NF) serving as the reference group for treated conditions. AT, anterior tubule; PT, posterior tubule; NF, normal food; BG, [specify compound name as defined in the main text].

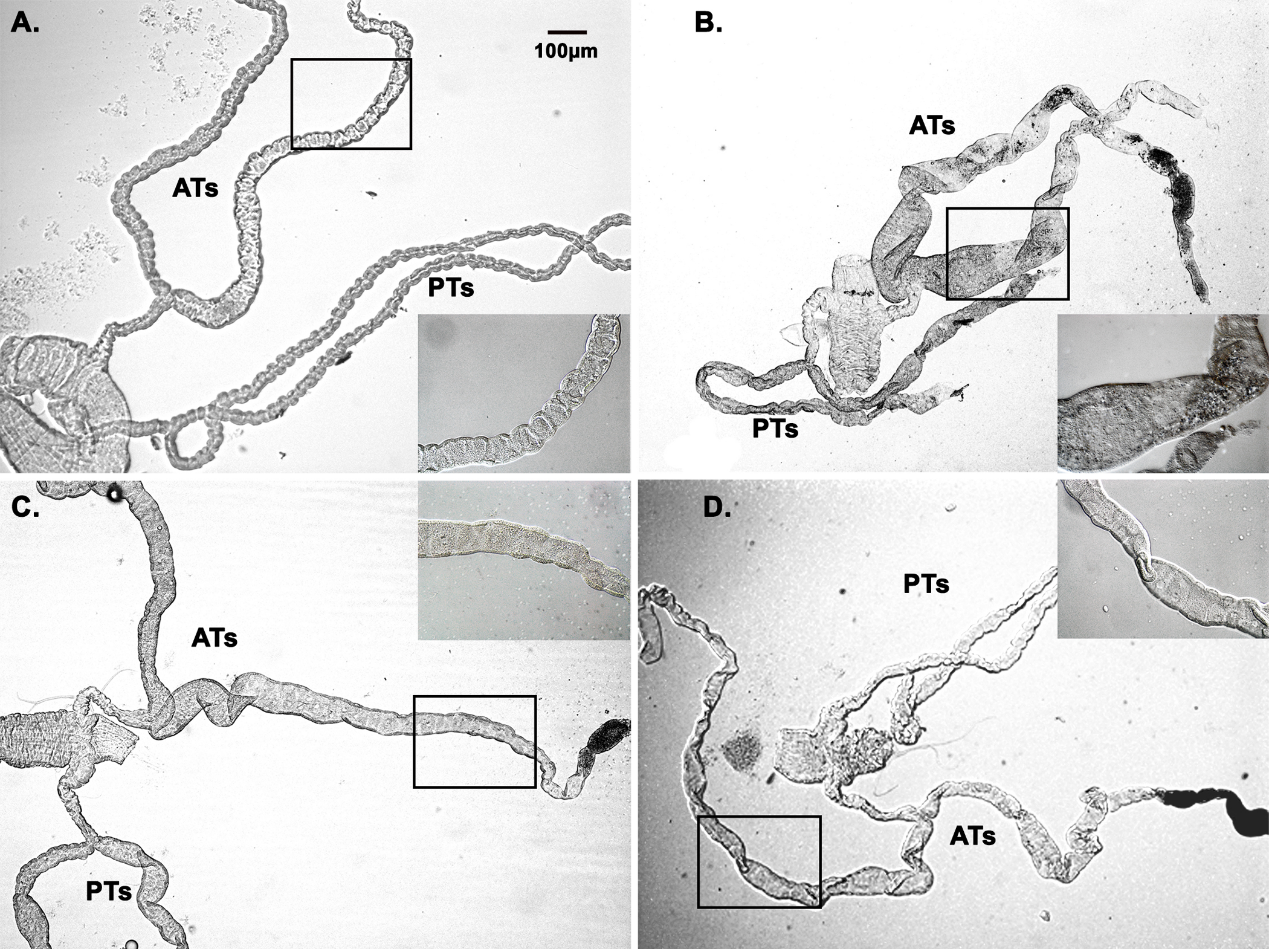

**Supplementary Image S6: MTs morphology upon BG supplementation.** Drice^∆2C8^, NF group (B) show highly dilated cystic MTs when compared to the wild type NF group (A). BG 0.25% (C) and BG 0.50% (D) supplementation restored the MTs morphology with reduction in tubule width and cyst formation in MTs. These are the DIC images of the MTs.

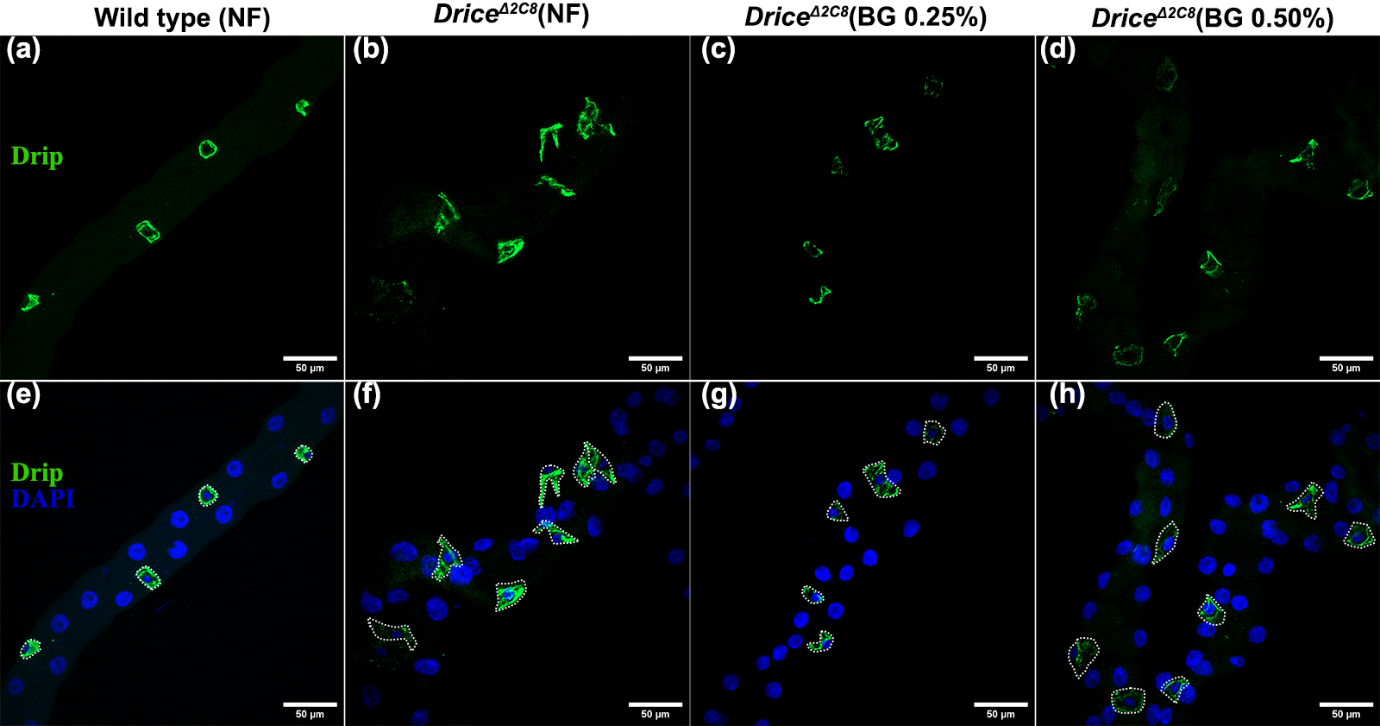

**Supplementary Image S7: BG Supplementation restores the SCs shap.** Drice^∆2C8^, NF group (b & f) show polygonal star shaped SCs when compared to the wild type NF group showing rectangular/cuboidal shaped SCs (a & e). BG 0.25% (c & g) and BG 0.50% (d & h) supplementation restored the SCs shape along with an increase in SCs number in MTs of Drice deletion mutants. Magnitude of the scale bar is 50 µm. Images are showing the confocal projection images of the MTs.

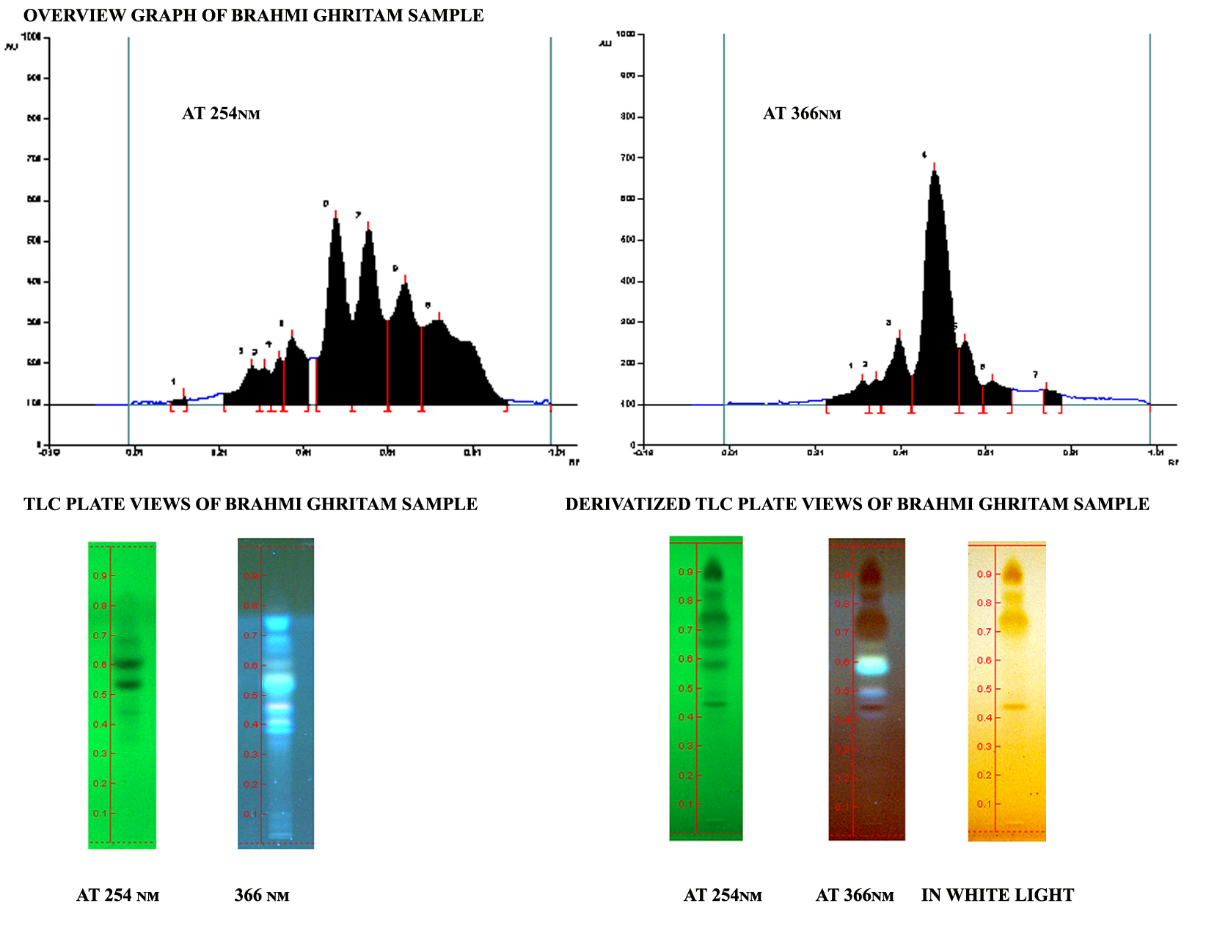

**Supplementary Image S8: HPLC profiling of the BG used for the formulation supplementation.**
